# CryptoCEN: A Co-Expression Network for *Cryptococcus neoformans* reveals novel proteins involved in DNA damage repair

**DOI:** 10.1101/2023.08.17.553567

**Authors:** Matthew J. O’Meara, Jackson R. Rapala, Connie B. Nichols, Christina Alexandre, R. Blake Billmyre, Jacob L Steenwyk, J. Andrew Alspaugh, Teresa R. O’Meara

## Abstract

Elucidating gene function is a major goal in biology, especially among non-model organisms. However, doing so is complicated by the fact that molecular conservation does not always mirror functional conservation, and that complex relationships among genes are responsible for encoding pathways and higher-order biological processes. Co-expression, a promising approach for predicting gene function, relies on the general principal that genes with similar expression patterns across multiple conditions will likely be involved in the same biological process. For *Cryptococcus neoformans,* a prevalent human fungal pathogen greatly diverged from model yeasts, approximately 60% of the predicted genes in the genome lack functional annotations. Here, we leveraged a large amount of publicly available transcriptomic data to generate a *C. neoformans* Co-Expression Network (CryptoCEN), successfully recapitulating known protein networks, predicting gene function, and enabling insights into the principles influencing co-expression. With 100% predictive accuracy, we used CryptoCEN to identify 13 new DNA damage response genes, underscoring the utility of guilt-by-association for determining gene function. Overall, co-expression is a powerful tool for uncovering gene function, and decreases the experimental tests needed to identify functions for currently under-annotated genes.

## Introduction

The availability of genome sequences from non-model organisms has greatly increased with the development of high-throughput sequencing techniques; however, insights into gene function have not kept pace. Without functional information, it is difficult to interpret the results of genome-scale experiments, hindering our ability to model and predict cellular function. There are several computational methods for predicting protein function, including machine-learning approaches based on sequence [1,2], structure [3], homology, literature co-reference, and integrated bioinformatics [4]. Co-expression, the coordinated regulation of two genes across various conditions, is observed among genes with similar function [5], and the principle of guilt-by-association has proven to be a successful approach for predicting gene function based on co-expression patterns [6–8]. Co-expression has consistently been a strong source for gene function prediction; for example, a global effort aiming to improve gene function prediction — the Critical Assessment of Function Annotation 3 challenge — found that expression-based approaches outperformed all others [9].

Fungal pathogens pose a growing threat to human welfare. *Cryptococcus neoformans*, an opportunistic human fungal pathogen that causes lethal meningitis in immunocompromised individuals if left untreated, was recently given the highest priority of public importance by the World Health Organization [10]. Of the major human fungal pathogens, *C. neoformans* is also notable for being a basidiomycete, having diverged from the ascomycete lineage, which contains the model yeast *Saccharomyces cerevisiae* and the majority of human fungal pathogens approximately 400 million years ago [11]. Much of the current gene annotation information comes from inferred orthology with *S. cerevisiae*, but only 17% of *C. neoformans* genes have *S. cerevisiae* ortholog annotations due to the large evolutionary distance between these organisms. Thus, nearly 60% of the genes in the *C. neoformans* genome remain “hypothetical” or “unspecified” and lack any computed or curated biological process Gene Ontology (GO) term information [12,13]. Additionally, there have been numerous examples where sequence conservation between *C. neoformans* proteins and model yeast failed to predict gene function [14–16].

As *C. neoformans* is less genetically tractable than model yeast, researchers have taken advantage of transcriptomic approaches to help understand how the cells are responding to a variety of environmental and genetic perturbations. For example, differential expression analysis helped identify multiple stages of *C. neoformans* infection of the host [17], including the extensive cell wall remodeling during host infection. Other studies have performed transcriptional profiling across mutant strains to identify regulatory relationships between transcription factors that control capsule formation, the key virulence factor in *C. neoformans* [18], or the response to environmental pH [19]. Previous investigators have used computational approaches to predict gene function in *C. neoformans* through the generation of CryptoNetV1 [20]. However, advances in transcriptomics, predictive algorithms, and structural modeling now allow for more rigorous strategies in assigning potential function to unannotated genes.

Here, we leveraged the extensive publicly available transcriptional profiling data generated by the *C. neoformans* research community to build a *C. neoformans* co-expression network (termed CryptoCEN) that captures multiple dimensions of genetic and cellular function. For example, CryptoCEN uncovered the capsule co-expression network that encompassed cell cycle processes, revealed a co-expression signature between ergosterol and translation, and identified a role for the Cdc42 and Cdc420 paralogs in kinetochore assembly. Importantly, we were also able to demonstrate the high predictive value of co-expression by prospectively identifying 13 new proteins involved in DNA damage responses, a set of pathways responsible for maintaining genome integrity that are thought to be promising targets for therapeutics [21]. Together, this demonstrates the functional utility of co-expression in understanding gene function.

## Results

### Generating CryptoCEN, a global co-expression network that captures genomic function

We generated a co-expression network for *C. neoformans* based on the general principles established in our previous implementation of a co-expression network for *C. albicans* using our CalCEN R package [7]. First, we collected RNA sequencing (RNAseq) data from the NCBI Sequence Read Archives (SRA). We chose studies that included at least 8 samples and filtered for those examining the *C. neoformans* H99 type strain and its derivatives, including KN99. This resulted in 1,524 runs across 34 studies, listed in **Table S1**. The conditions for these experiments included a wide range of environmental perturbations, such as differences in nutrient source, cell cycle, chemical perturbations, and genetic mutation [18,19,22–27]. For consistency, the raw reads from each study were re-mapped to the 9,189 genes from FungiDB release-49 of the *C. neoformans* H99 reference transcriptome using RSEM with bowtie2 [28,29], thus ensuring uniformity in analysis and data processing. Similar to our work in *C. albicans,* to ensure sufficient coverage, we removed runs where greater than 50% of the genes had zero expression, yielding 1,523 samples in total (**Table S1**). We used fragments per kilobase of transcript per million mapped reads (FPKM) as the estimated expression for each gene under each condition, and then used Spearman rank correlation to measure the correlation between gene expression profiles with the EGAD R package [7,30]. Rather than pooling all expression profiles for all studies, we built separate co-expression networks for each study and combined the average across all networks: for studies designed to explore specific biology, the within-study expression variation may be more meaningful than the between-study variation. For each pair of *C. neoformans* genes, we then generated a value between 0 and 1 to represent the rank of co-expression among all pairs of genes **(Data 1)**.

We then used UMAP dimensionality reduction and Louvain clustering methods [31,32] to visualize the overall *C. neoformans* co-expression network (CryptoCEN); this approach aims to keep co-expressed genes clustered together (**Figure 1A**). We identified 22 clusters (**Table S2**) and used GO-term enrichment analysis using Fisher’s exact test run through FungiDB of biological process to predict functional signatures for each group. There was a clear cluster for cell cycle proteins and DNA replication (cluster 10) and proteins involved in carbohydrate transport (cluster 13). We also observed sub-structures within each cluster, such as in cluster 6, where there were sub-clusters for the large and small ribosome proteins and for oxidative response proteins (**Figure 1B).** Additionally, in cluster 16, which was enriched for proteins involved in respiration, we could see a distinct sub-cluster for those proteins encoded within the mitochondrial genome (**Figure 1C).** Recapitulating known functional groups of genes demonstrate the efficacy of our approach. However, we also identified 8/22 clusters for which there was no enrichment for any biological process GO term annotation, highlighting the lack of functional information for many genes in the *C. neoformans* genome.

**Figure 1:**
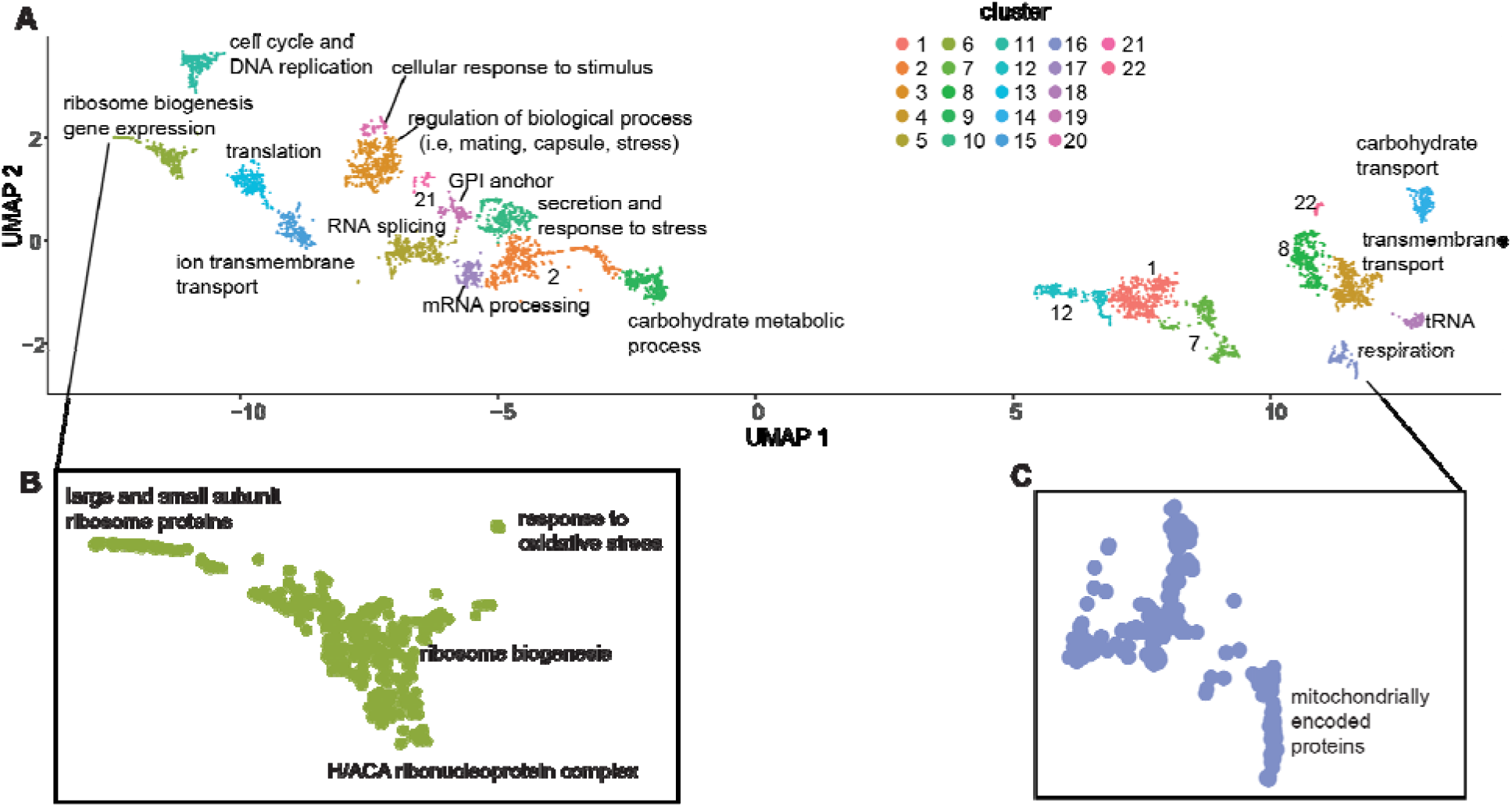
*Cryptococcus* Co-Expression analysis identifies both known and unknown gene clusters. A) Embedding of the *C. neoformans* Co-Expression Network (CryptoCEN) using UMAP for dimensionality reduction. Twenty-two clusters were identified using Louvain modularity clustering. The genes in each cluster were analyzed for biological process GO term enrichment using Fisher’s exact test through FungiDB, and the most significantly enriched specific term after Bonferroni correction for multiple testing was used for labelling. Clusters without significant GO term enrichment after multiple hypothesis testing correction were labeled only by cluster number. B) Genes in cluster 6 sub-clusters were analyzed for both biological process and cellular component GO term enrichment using Fisher’s exact test through FungiDB. C) Genes in cluster 16 sub-clusters were analyzed for both biological process and cellular component GO term enrichment using Fisher’s exact test through FungiDB.

### Benchmarking CryptoCEN via Retrospective Gene Function Analysis

There are 8,334 genes currently identified in the *C. neoformans* genome, and 2754 of them are uncharacterized with no functional annotation across homology or gene function prediction tools **(Figure 2A).** A goal of the co-expression network is to predict gene function for these underannotated genes. Therefore, we first benchmarked the ability of the CryptoCEN to retrospectively predict all of the current *C. neoformans* GO term annotations without the NOT qualifier collected from FungiDB [33]. Since GO terms form an ontological hierarchy, we propagated annotations from more specific to more general terms and then filtered for terms having between 20 and 1,000 annotations. For this benchmarking, we used guilt-by-association, where the strength of the GO term prediction is based on the fraction of neighbors (or weighted sum for a network where edges are weighted e.g., by co-expression score) in the network with a given term. We then compared our predictions with the current set of *C. neoformans* predictions using the area under the receiver operating characteristic curve (AUROC) score and averaging over a 10-fold cross-validation. True-positive results were defined when the CryptoCEN predicted GO term matched the known GO term from FungiDB and the AUROC score provided an unbiased measure of enrichment and network quality. Using this dataset, we found that the CryptoCEN network had a neighbor-voting AUROC of 0.74 (+/- 0.093) (**Figure 2B**), suggesting that CryptoCEN was not making predictions based on multifunctional genes [7,34].

**Figure 2:**
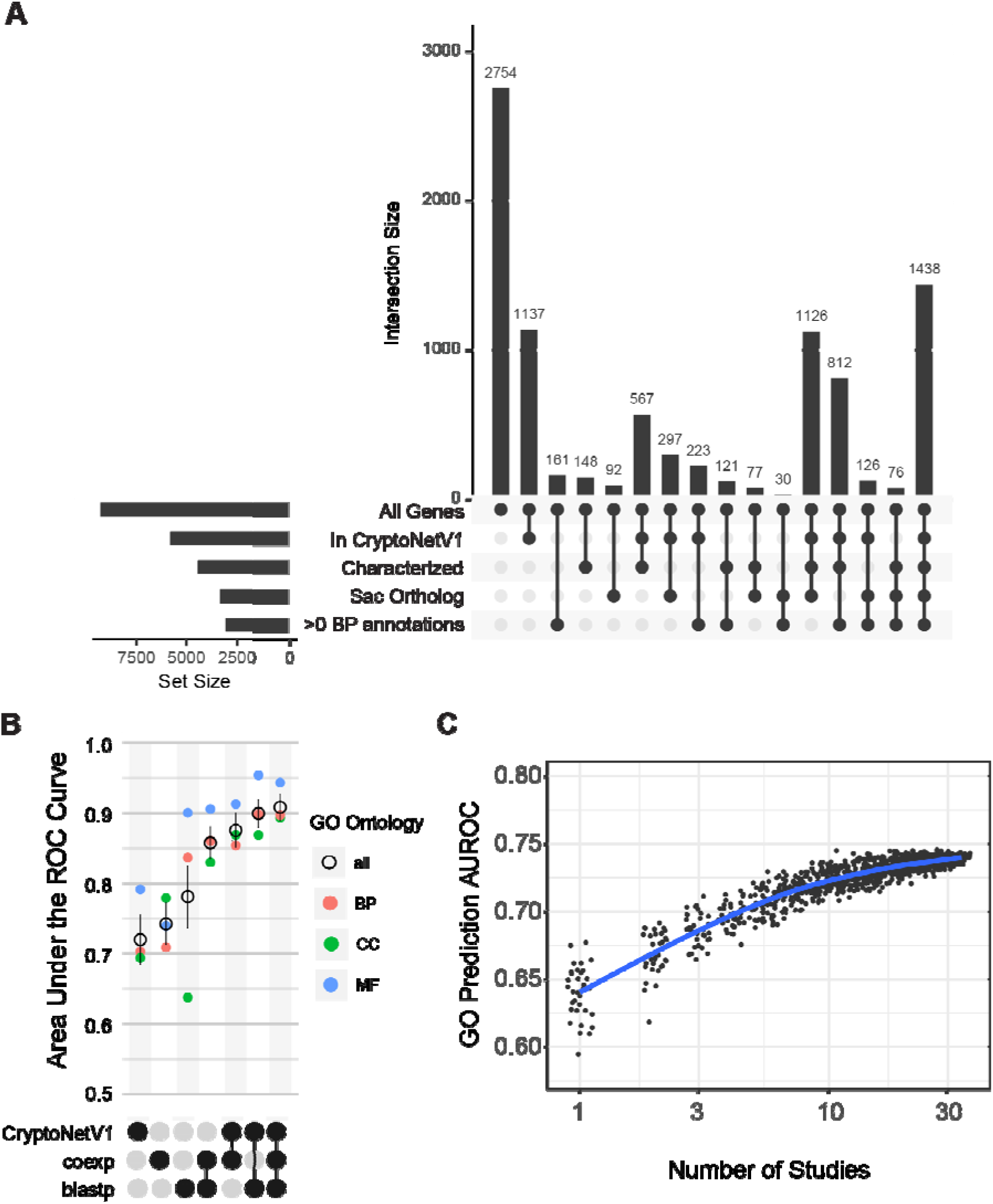
CryptoCEN can retrospectively predict biological process GO terms. A) The *C. neoformans* genome contains many unannotated genes. UpSet plot of gene annotation information from different sources. Each bar in the upper region shows the number of gene nodes in the intersection of the set of databases indicated by the rows with filled circles in the lower region. The first column represents genes that are not included in any of the current annotation databases, including orthology to the model yeast *S. cerevisiae*. B) Combining CryptoCEN with other sources of information increases retrospective prediction accuracy. UpSet plot of the retrospective prediction accuracy, as determined by the neighbor voting guilt-by-association (GBA) area under the ROC curve (AUROC). AUROCs values range between 0.5 for random predictor and 1 for a perfect predictor. As data sources are combined, the prediction accuracy increases. Each annotated GO term is colored by ontology biological process (BP), cellular component (CC), or molecular function (MF). C) Prediction accuracy increases as the number of studies included in the network increases. Mean neighbor voting GBA performance for the CryptoCEN built over random subsets of RNAseq studies. The blue curve represents a mean of a nonparametric locally estimated scatterplot scattering (LOESS) fit.

In comparison, using information from orthologous systems, such as BLAST or CryptoNetV1 [20], resulted in AUROCs of 0.78 (+/- 0.14) and 0.72 (+/- 0.11). Using orthologous physical or genetic associations for *S. cerevisiae* had a predictive power roughly on par with random chance (AUROCs of 0.56 ± 0.095 and 0.55 ± 0.87, respectively) (**Figure S1**). In contrast with BLAST or CryptoNetV1, which had better relative prediction over biological process and molecular function, the CryptoCEN network had a relatively better prediction for the cellular component terms. To quantify if orthologous systems are complementary, we evaluated AUROC over all combinations of networks by summing over edge weights. The predictive performance of CryptoCEN in combination with BLAST and CryptoNetV1 offered a small but measurable increase compared to the combined predictive power of the two established models. For both BLAST and CryptoNetV1, adding CryptoCEN substantially increased performance to AUROCs of 0.86 (+/- 0.071) and 0.88 (+/- 0.076) respectively, and all three combined lead to the highest overall performance of 0.91 (+/- 0.06), compared with 0.90 (+/- 0.64) for BLAST and CryptoNetV1 only. Using all three networks specifically had a benefit for biological process and cellular component prediction **(Figure 2B).** This demonstrates that CryptoCEN captures information undetected in previous prediction tools and that adding RNAseq-based co-expression can increase the quality of gene function predictions, at least retrospectively.

A potential limitation of CryptoCEN is that the transcriptome space has not been fully explored, and additional RNAseq studies may be needed to break spurious correlated expression by measuring under new environmental contexts. To address this question, we assessed how the predictive accuracy of the network changes as we remove studies from the network. For each k in the range [1, 33], we sampled different combinations of studies of size k and estimated their ability to predict gene function annotations. We found that as we increased the studies from 1-10, there was a rapid rise in average performance from ∼0.64 to ∼0.71. After this point, performance increases steadily to 0.74 when approaching the full set of 34 studies (**Fig 2C**). This suggests that we have sufficient studies of the transcriptome to build a robust co-expression network, but that the accuracy will improve with the integration of additional studies. Although the number of RNAseq studies needed to generate a co-expression network will vary based on genetic background and the fraction of transcriptome space represented, this finding may also help inform the experimental design to generate co-expression networks in other organisms.

### Evolutionary constraints can inform co-expression

Among proteins that form physical interactions, we hypothesized that those participating in obligate complexes may have stronger selective pressure for co-expression. For example, if all the sub-units need to be expressed at stoichiometric levels for the complex be functional, then the metabolic cost of expressing isolated subunits is wasted, leading to a selective pressure for co-expression [35]. Moreover, incorrect expression outside of stoichiometric ratios can generate proteotoxic stress, as the uncomplexed subunits are actively detrimental to the cell, as demonstrated during cases of aneuploidy [36].

Given that there is not a well-curated list of complexes in *C. neoformans*, we leveraged the better annotated complexes in *S. cerevisiae* and used sequence homology to infer candidate complexes. To support this strategy, we reasoned that complexes that are highly evolutionarily conserved may have robust cooperative function. We collected *S. cerevisiae* complexes from EBI (2021-10-13) and filtered out sub-units that were nucleic acid, small molecule, mitochondrially encoded, or unrecognized, yielding 616 multi-subunit complexes. Of these, 408 had at least 2 distinct subunits with a one-to-one ortholog in *C. neoformans* identified by OrthoMCL [37]. Within these complexes, there are 13,950 pair-wise co-complex interactions of *C. neoformans* proteins **(Table S3)**. Of these, over half (7,142) are interactions within the 17 annotated ribosomal or ribonucleic protein complexes, with an average co-expression score of 0.84. The other half of the candidate interactions (6,808) had an average co-expression score of 0.75 **(Figure 3A)**. This finding supports the hypothesis that there is selective pressure to co-express members of protein complexes.

**Figure 3:**
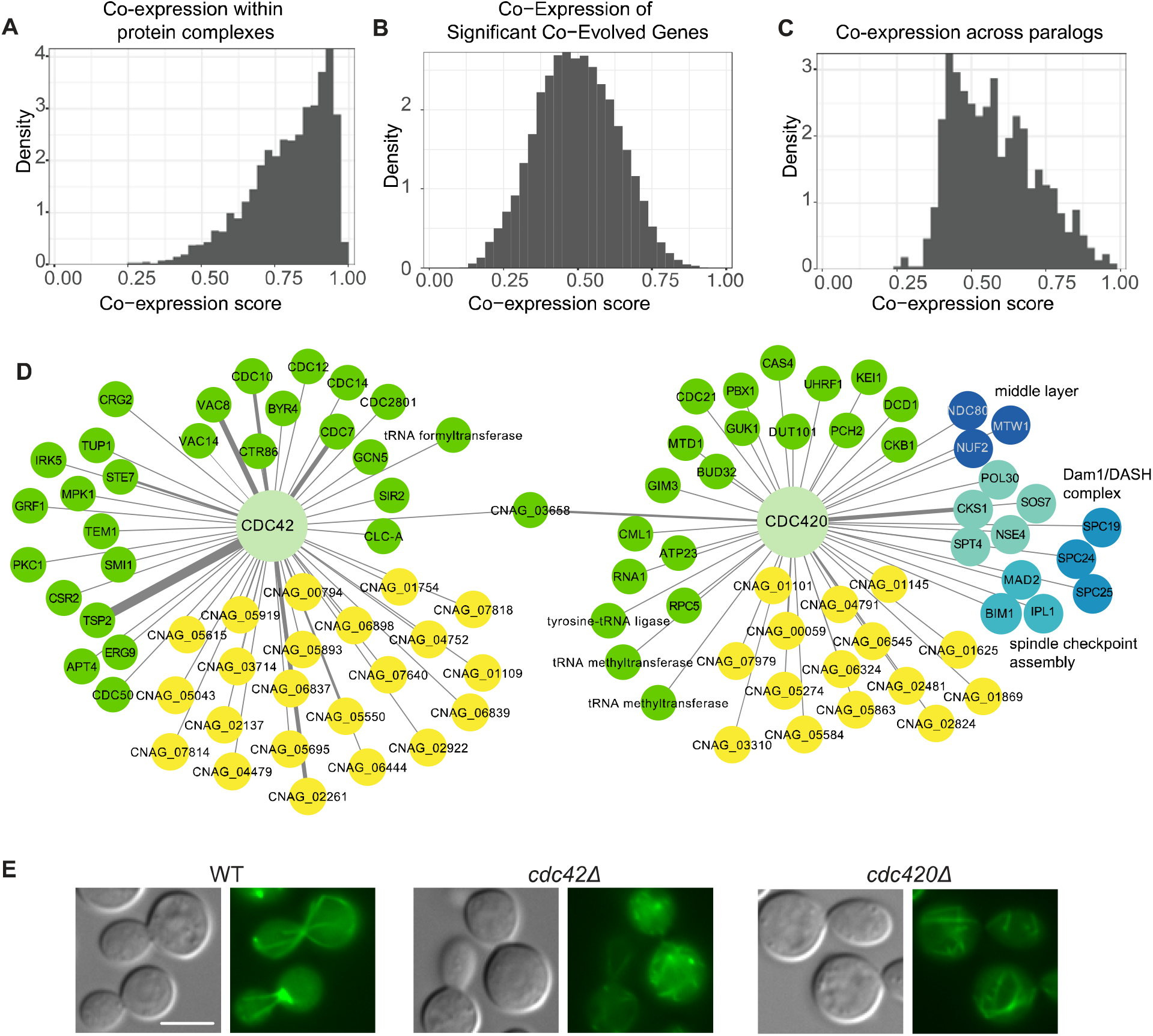
Evolutionary constraints inform co-expression analyses. Distribution of co-expression scores for *C. neoformans* gene pairs across different types of evolutionary conservation. A) *C. neoformans* gene pairs with orthology to *S. cerevisiae* gene pairs that encode for proteins that are members of the same complex (13,950 pairs over 1,304 genes, coexp score mean: 0.80, IRQ50: [0.71, 0.90]), B) Significantly co-evolving gene pairs (140,592 pairs over 4,269 genes, coexp score mean: 0.50, interquartile-range at 50% (IRQ50): [0.40, 0.60]), and C) Paralogous gene pairs (1,056 pairs over 550 genes, coexp score mean: 0.58, IRQ50: [0.46, 0.67]). D) The co-expressed partners of the duplicated genes Cdc42 and Cdc420 were compared and visualized in Cytoscape. Kinetochore genes are highlighted in shades of blue. Unannotated proteins are highlighted in yellow. Width of the lines indicates co-expression score. E) Tubulin is altered in the *cdc420*Δ and *cdc42*Δ mutant strains compared with the H99 wildtype. Tubulin was visualized by fusion of α-tubulin with GFP. Cells were incubated in liquid YPD at 30 °C before imaging. Images taken at 40X magnification, scale = 5 microns.

A recently developed approach to predict function is to use co-evolutionary networks; beyond being part of the same complex, co-evolution is based on the principle that functionally related genes will show similar rates of evolution across speciation events [38]. To calculate this, the evolutionary rate values are estimated for each branch of an orthologous gene’s phylogeny and compared with each other gene to determine the rate of co-evolution [38]. We hypothesized that combining a co-evolutionary network would increase the accuracy of the co-expression network. Therefore, we generated a co-evolutionary network for *C. neoformans* using 15 related species (**Figure S2B**), with the caveat that the species tree of related organisms is much sparser for *C. neoformans* than for *S. cerevisiae,* making this analysis less robust that the established *S. cerevisiae* co-evolution information [38]. We then examined whether combining the co-evolutionary network would increase the accuracy of the co-expression network. Here, we subset the analysis to those genes with clear ortholog groups (**Table S4**). However, we identified very little correlation between co-expression and co-evolution networks (**Figure 3B**), limiting our ability to integrate this information. This can be potentially attributed to the low density of closely related species genome sequences, which limits our ability to identify co-evolution signatures.

It is also possible to examine the evolutionary pressures on co-expression in the context of gene duplication. Gene duplication provides an opportunity for either shared functionalization, as in the case of the histone proteins, or neofunctionalization and sub-functionalization [39,40]. Among genes with clear ortholog groups (**Table S4**), we examined the co-expression of each protein with every other protein in each orthogroup and plotted the distribution (**Figure 3C**). Given that the co-expression scores are ranked, a random distribution should be flat. However, we observed that the co-expression is less than random, suggesting that there is pressure to be differentially transcribed from other genes within an orthogroup. To characterize the distribution, we fit the distribution with a skew-normal curve, and the alpha parameter is significantly above zero (4.4 with a standard deviation of 0.56), indicating the extent of the skew. The outlier with high co-expression between orthologous genes were the H2A and H2B histone genes. This bias towards a shift in expression between paralogs is consistent with previous results in multiple systems, where changes in transcription are required for evolutionary divergence in function [41–43].

In *C. neoformans,* we have also observed the duplication of entire signaling cascades, as opposed to single gene duplications [44]. One notable example is the duplication of Ras1/Ras2, Cdc42/Cdc420 and Rac1/Rac2 proteins [44–46], with each paralog supporting specific cellular functions. For example, Cdc42 serves as a critical regulator of thermotolerance and virulence [46,47], while Cdc420 plays a minor role in the expression of virulence-associated phenotypes under basal conditions [48]. Importantly, Cdc42 is critical for septin localization, with Cdc10 completely mis-localized in the *cdc42*Δ mutant, likely explaining similar defects in thermotolerance in the *cdc42*Δ and septin mutant strains [48].

We analyzed the co-expression partners for the paralogous Cdc42 and Cdc420 proteins, with visualization using Cytoscape, where nodes represent genes and edges connect two co-expressed genes. From this, we identified almost entirely distinct networks for the two paralogs, with only the CNAG_03658 hypothetical protein co-expressed with both Cdc42 and Cdc420. Consistent with prior genetic and physiological data, Cdc42 was highly co-expressed with multiple septin proteins, including Cdc10 **(Figure 3D)**. In contrast, Cdc420 was co-expressed with 10 kinetochore or spindle pole body proteins **(Figure 3D),** including parts of the middle layer, outer layer (Dam1/DASH), and spindle assembly checkpoint [49–52]. Previous work on physical interaction partners for kinetochore proteins revealed that Spc25 interacts with Cdc420 [53], giving additional evidence for the predicted relationship between Cdc420 and kinetochores.

Therefore, to more directly test whether a loss-of-function mutation of Cdc420 would alter kinetochore function, we examined microtubule and nuclear dynamics in strains with either fluorescently-tagged α-tubulin (Tub1) or histone H4 proteins [49] in the WT, *cdc42*Δ, and *cdc420*Δ strains. The H4-Gfp protein highlighted a similarly well-defined nucleus in each strain and each growth condition **(Figure S2C)**. In the wild-type strain, expression of the Tub1-Gfp fusion protein demonstrated clear and elongated microtubules in cells undergoing cell division **(Figure 3E)**. In contrast, both the *cdc42*Δ and *cdc420*Δ strains showed more cells with punctate rather than polymerized GFP signal, consistent with a relative defect in α-tubulin polymerization into microtubules **(Figure 3E)**. Moreover, all strains showed similar growth in response to the microtubule destabilizing compound nocodazole (**Figure S2D**). These data support the predicted association between kinetochore function and both the Cdc420/Cdc42 proteins, and they also suggest that there is not a specific defect in microtubule assembly in the *cdc420*Δ compared with the *cdc42*Δ mutant strain. It is possible that the large set of uncharacterized genes (**Figure 3D)** that are co-expressed with *CDC42* and *CDC420* have roles in kinetochore or microtubule assembly, but the lack of current annotation prevents us from making those connections.

### Virulence factor retrospective cluster analysis

To understand how the co-expression network can identify new genes for a given function, we first retrospectively explored the network localized to the genes involved in well-studied functions.

The major virulence factor of *C. neoformans* is the polysaccharide capsule [54]. To seed the network we used a set of known capsule biosynthetic genes [54], identified all the first neighbors in the network with a co-expression score higher than 0.8, and then selected those with at least 5 co-expression edges to other genes in the set for visualization using Cytoscape [55] (**Figure 4A**). Interestingly, this analysis revealed that although Cap59, Cap60, Cap64 and Cas35 are highly interconnected and co-expressed, there are other capsule biosynthetic genes, such as Uge1 and Cas33 that are not directly co-expressed with any other capsule gene. Instead, they remain connected to the network through intermediaries. Given the importance of condition-dependent adaptations in other cell surface features to promote capsule attachment, we also saw co-enrichment for cell wall and membrane biosynthesis genes in this capsule gene cluster. The other major signature in the remaining genes was for proteins involved in cell cycle and DNA maintenance. This connection is consistent with previous literature on cell cycle and capsule in *C. neoformans* [16,56]. Notably, the only transcriptional regulators that appeared to show co-expression with capsule genes were orthologues to Ctk2 and the uncharacterized transcription factor Fcz4, despite the known importance of many other specific transcription factors in regulating capsule [15,57–59]. This complements the study from Kim et al., which focused on the capsule regulatory genes in CryptoNet and observed connections between the regulatory cascades but not the capsule biosynthetic genes [20]. Potentially, this is due to the regulation of signaling cascades at the post-translational level, rather than the transcript level.

**Figure 4:**
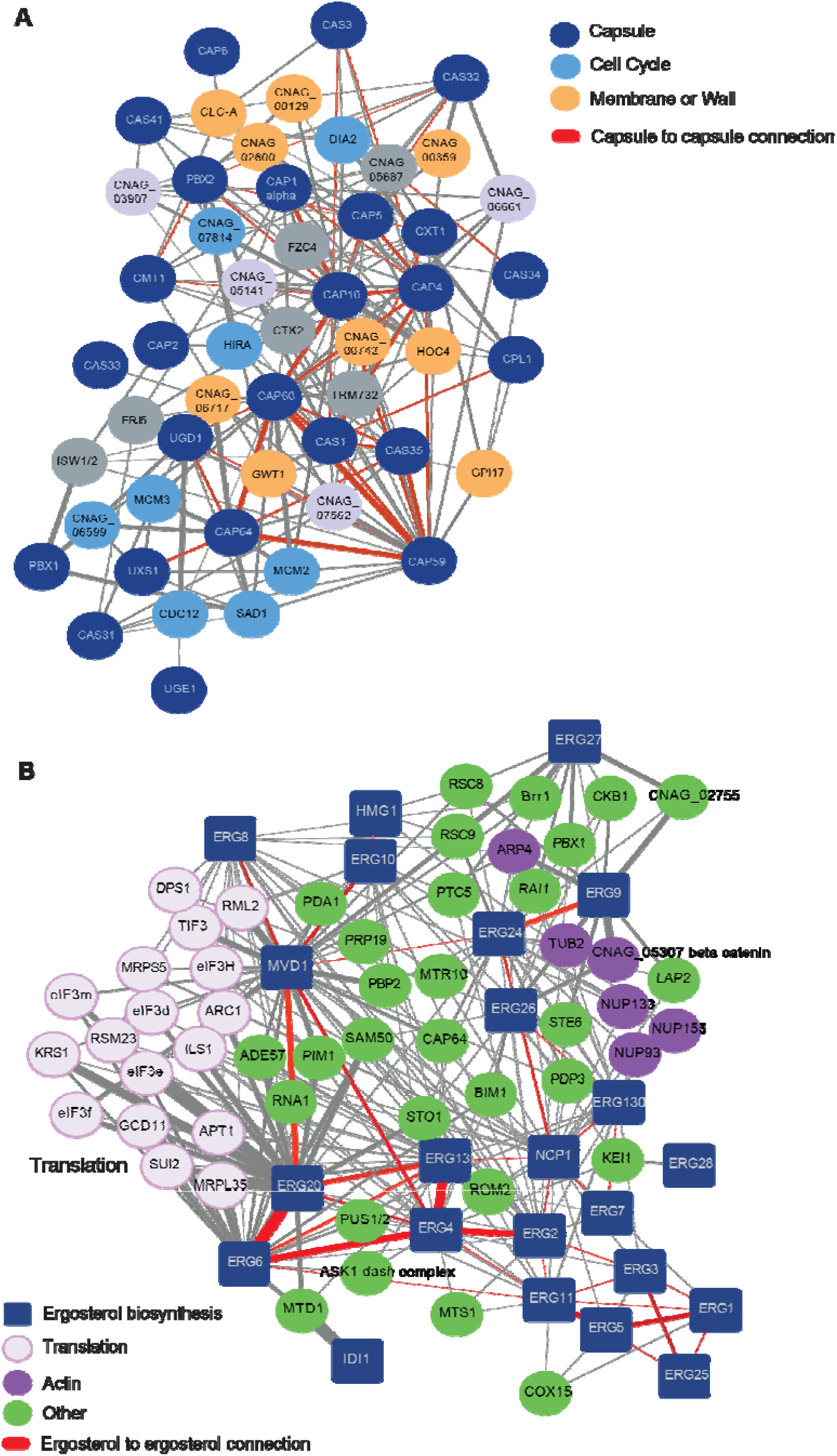
CryptoCEN can recapitulate core biological processes in *C. neoformans*. A) A co-expression network for capsule was generated by starting with genes known to be involved in capsule biosynthetic genes. All co-expressed partners with a score > 0.8 and at least 5 co-expression edges were visualized in Cytoscape. Specific functional classes are highlighted with different colors. Edge width corresponds to degree of co-expression. B) A co-expression network for ergosterol biosynthesis was started with the known ergosterol biosynthetic genes and all co-expression partners that showed >0.75 co-expression score and interaction with >3 ergosterol biosynthetic genes. Specific functional classes are highlighted with different colors. Edge width corresponds to degree of co-expression.

As a second test case, we examined the ergosterol biosynthetic cascade, as ergosterol and its biosynthesis are important antifungal drug targets [60]. We seeded the network with 23 orthologs of the known ergosterol biosynthetic machinery and filtered for the first neighbors with at least three co-expression edges with ergosterol biosynthesis proteins and a score of at least 0.75 **(Figure 4B)**. This resulted in a densely connected network, but with a somewhat surprising structure—the ergosterol genes were not organized in biosynthetic order. For example, Erg1, Erg3, Erg5, and Erg25 were highly interconnected despite operating in different parts of the ergosterol biosynthesis pathway. We also observed many proteins involved in translation with strong co-expression with the ergosterol genes, especially Erg20 and Erg6.

Overall, we were able to consistently replicate known networks through CryptoCEN, and potentially identify new signatures associated with core biological processes. This suggests there is utility in using co-expression to explore gene function in *C. neoformans*.

### Identification of additional genes involved in DNA repair, including novel uncharacterized proteins

DNA repair is an essential and highly conserved function in cells that ensures genome stability. From a biomedical perspective, SNPs in mismatch repair genes have been linked with hypermutator phenotypes in *C. neoformans,* allowing for increased drug resistance and virulence [61–63]. From an evolutionary perspective, comparative genomic analysis has revealed gene presence/absence variation among canonical DNA repair genes in other microeukaryotes [64,65], but the discovery of novel DNA repair genes has been lacking. In *C. neoformans*, the full set of DNA repair genes is unknown. We hypothesized that genes coexpressed with those known to contribute to DNA repair—34 ERCC, MLH, MSH and RAD proteins identified by Ashton and colleagues [66]—would allow us to identify additional uncharacterized proteins involved in this core biological process.

To do so, we filtered for co-expression scores > 0.8 and at least four co-expression edges with known DNA repair genes, and then used this set to generate a network that we visualized in Cytoscape (**Figure 5A**). This revealed a dense network of mismatch repair genes centered around *MSH2* and *MSH6*, and a sparser network with more of the ERCC excision-repair genes. The initial seed set of DNA damage response genes (purple) was surrounded by genes with known functions in DNA damage and repair, including the *C. neoformans-*specific radiation resistance transcription factor, Bdr1 [67]. The centrality of some of the cell cycle and DNA replication genes in this network, including Bud14, Kel1, Pol3 and Rig2/Dbp11, highlights the strong association between DNA repair and cell cycle processes.

**Figure 5:**
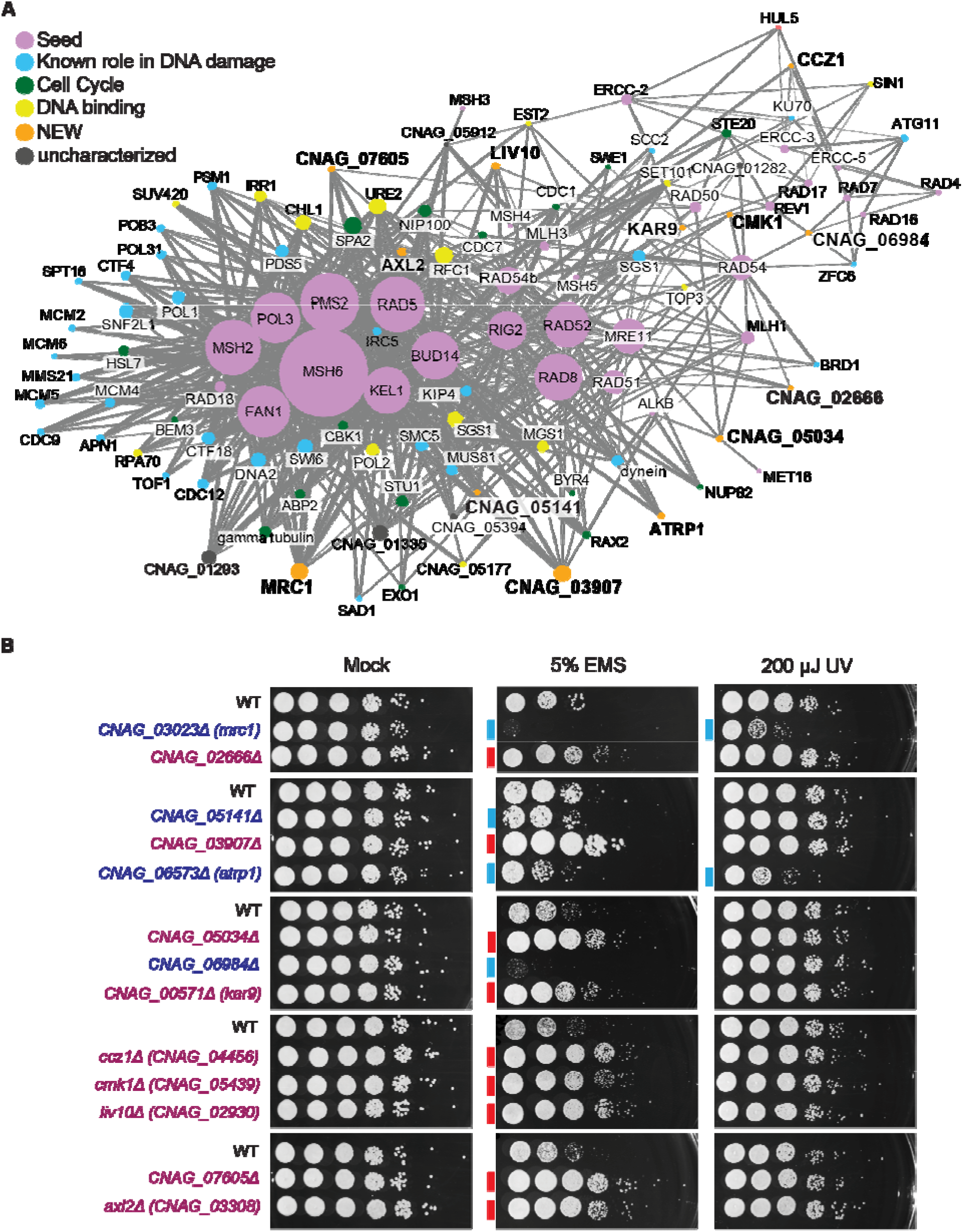
Identification of new proteins involved in DNA damage responses. A) A co-expression network for DNA damage was started with 34 known genes involved in DNA repair, and all co-expression partners that showed > 0.8 co-expression score and interaction with at least 5 of the known DNA repair genes. Specific functional classes are highlighted with different colors. Edge width corresponds to degree of co-expression. B) Identification of novel genes are involved in DNA damage responses. The indicated strains were grown overnight at 30°C in liquid YPD medium, and the 10-fold serially diluted cells were spotted onto YPD agar. For UV damage, the plates were immediately subjected to 200 µJ UV. For EMS, the cells were incubated in 100 µM EMS for 1 hr before serial dilution and plating. The plates were incubated at 30°C and imaged after 2 days.

Beyond the known proteins in this DNA damage response network, we identified many highly co-expressed hypothetical proteins that we hypothesized may play a role in DNA repair, including proteins of unknown function as well as those with known roles in other aspects of cell biology. Therefore, we tested 13 available deletion mutants of these strains for their responses to either ethyl methanesulfonate (EMS) or UV radiation, as a measure of two different DNA damage response pathways—DNA alkylation and DNA lesions (**Table S6**). Notably, all 13 of the mutants tested had a phenotype on EMS, including 9 resistant and 4 sensitive strains **(Figure 5B)**. Of the four EMS-sensitive strains, two were also sensitive to UV damage.

To determine whether the predictive accuracy was above baseline, we tested a set of random mutants for their phenotypes on DNA damaging agents. A potential complication is that DNA replication and damage responses are amongst the most highly co-expressed genes in CryptoCEN. Therefore, we chose 12 genes that showed a similar rank of co-expression as our matched control set for determining the baseline rate of phenotypes in response to DNA damaging agents. Here, only 4 of the genes showed a phenotype **(Figure S2)**, with two strains showing sensitivity to EMS and two showing slight resistance. Therefore, CryptoCEN has high prospective predictive accuracy for gene function annotation.

One of the UV-sensitive mutants was *MRC1*/CNAG_03023, a hypothetical protein with an MRC1 (mediator of replication checkpoint)-like domain with 13 co-expressed partners in the network. When we performed the reciprocal co-expression analysis using *MRC1* as the seed, in addition to the 13 previously identified partners, we identified an additional 13 genes with roles in chromatin binding, DNA damage responses, or cell cycle **(Table S6)**. The presence of the MRC1 domain in the CNAG_03023 sequence suggested a putative function for CNAG_03023 as a checkpoint protein that would be required for the response to multiple types of DNA damage.

The other UV-sensitive strain was the deletion for CNAG_06573, a basidiomycete-specific hypothetical protein with 5 co-expressed partners. This protein did not have any conserved domains, and although reciprocal co-expression analysis identified a further 18 proteins involved in DNA damage, cell cycle, and DNA binding, there was not a clear signature of function beyond this larger category **(Table S6)**. Therefore, we turned to structural homology, using AlphaFold Monomer v2.0 2022-11-01[68] to give a predicted structure to use as an input to FoldSeek [69]. The well-structured N-terminal domain of CNAG_06573 (residues ∼700-1200) had homology with the CATH Superfamily [70] Leucine-rich Repeat Variant (1.25.10.10), which contains DNA repair protein rad26 (rad26) from *S. pombe* (sequence identify of 11.4 and E-value of 2.5e-4), and the ATR-interacting protein (ATRIP) from *H. sapiens* (sequence identity of 11.2 and E-value of 2.1e-2) [71,72]. ATRIP domains recognize the Replication Protein A complex associated with single stranded DNA to facilitate DNA damage response [71,72]. Therefore, although CNAG_06573 does not share sequence similarity with these known DNA-damage response proteins, the structural homology suggests that CNAG_06573 may act as an ATRIP in *C. neoformans.* Based on this, we named CNAG_06573 as *ATRP1*.

The *CNAG_05141*Δ mutant had minor sensitivity to EMS compared to the wild-type control, and was not sensitive to UV damage. Based on AlphaFold and FoldSeek analysis, this protein contained a predicted EamA-like transporter domain [73] and had structural homology to the Nipal2 and Nipa1 transporter proteins. Potentially, loss of this transporter may lead to a higher intracellular concentration of EMS resulting in increased cell death. Notably, this gene was also co-expressed with multiple capsule biosynthetic genes.

The *CNAG_00571*Δ mutant also displayed an EMS-resistant phenotype. This gene encodes a hypothetical protein with a predicted karyogamy protein 9 (KAR9) domain. Kar9 facilitates nuclear congression during karyogamy in *S. cerevisiae* [74]. Previous work on the karyogamy machinery in *C. neoformans,* however, had indicated that an ortholog to *S. cerevisiae KAR9* (*ScKAR9*) was not present in *C. neoformans* [75]. These searches were based on sequence homology and synteny, and CNAG_00571 is not syntenic to *ScKAR9* and does not share major sequence homology, despite the predicted Kar9 domain. However, yeast are known to evolve rapidly, resulting in low syntenic conservation [76], and remote homology may be difficult to detect [77]. Using a structural homology-based approach, we found that the predicted structure of CNAG_00571 shared with homology with the CATH superfamily 1.20.58.70, which is enriched in syntaxin proteins which are known to facilitate membrane-membrane fusion events within the cell. However, the lack of sequence homology with *ScKAR9* and the placement of CNAG_00571 in a Tremellales-specific ortholog group (OG6-532064) indicates that this gene does not share an evolutionary history with *ScKAR9*. Despite this, we hypothesize that CNAG_00571 is indeed functioning as a Kar9 protein during karyogamy by facilitating nuclear-nuclear fusion in *C. neoformans*, and thus we propose naming CNAG_00571 as *KAR9*.

We attempted a similar structural-based approach for CNAG_06984, which showed specific hypersensitivity to EMS. This protein may be involved in double-stranded break repair, based on the specific hypersensitivity phenotype [78]. However, neither structure nor sequence-based approaches yielded any related proteins with high confidence. Furthermore, three genes specific to the Tremella lineage—CNAG_02666, CNAG_07605, CNAG_02930, whose mutants were all resistant to EMS—had low-quality structural predictions, likely due to a lack of the required training data.

Beyond the uncharacterized proteins, we also identified two genes that had previously been implicated in different cellular pathways, but still showed an EMS-resistant phenotype. Ccz1 (*CNAG_04456*) is a guanidine exchange factor (GEF) that forms an active complex with Mon1 (*CNAG_00971*) [79], although these two genes are not co-expressed in *C. neoformans*. In *S. cerevisiae*, this Ccz1-Mon1 complex activates Rab7, which is involved in intracellular trafficking to the lysosome, including trafficking of the autophagosome to the lysosome [80–82]. Autophagy is known to be induced by DNA damage, where the inhibition of autophagy can sensitize cancer cells to DNA damage [83–85]. Therefore, we hypothesize that the loss of *CCZ1* in *C. neoformans* leads to a higher overall level of autophagy, which may provide a protective effect against DNA damage. GO term enrichment of *CCZ1* co-expression partners also showed enrichment for macroautophagy (p =1.87e-4).

The other previously annotated gene implicated in DNA repair using CryptoCEN is *CMK1*, which is more resistant to EMS. This gene encodes a calmodulin-dependent kinase (CaMK) and serves as an effector of the calcium-calcineurin signaling pathway, an important component of fungal stress responses [86–88]. Previous studies have shown that this pathway can play a role in cell cycle control in *S. cerevisiae* and other organisms [89], with a function at the G_2_/M checkpoint [90–92]. In *C. neoformans*, loss of *CMK1* may inhibit the mutant from the appropriate stress-induced cell cycle arrest, leading to increased growth in the presence of the cellular stress. GO term enrichment of the *CMK1* co-expression partners shows enrichment for DNA recombination, metal ion transport, and negative regulation of exit from mitosis (**Table S6)**. Together, this analysis of the DNA damage co-expression network identified multiple new proteins involved in DNA damage response in *C. neoformans,* including proteins without sequence or structural homology to known DNA damage response proteins.

## Discussion

*Cryptococcus neoformans,* despite being a deadly human fungal pathogen and the cause of mortality for nearly 200,000 people per year [93], has a genome that is vastly under-annotated. This lack of functional annotation information makes it difficult to interpret or identify genetic signatures or shed light on genotype-phenotype associations. Co-expression analysis provides a platform for predicting gene function, thus potentially decreasing the experiments needed to define the function of a gene. Recently, we generated a co-expression network for the model fungal pathogen *Candida albicans,* based on available RNAseq data from the reference strain SC5314, which allowed us to predict the function of gene as a cell cycle chaperone protein [7]. Co-expression across diverse clinical isolates of *C. albicans* was able to identify regulators of morphogenesis and virulence [94]. These, and others, demonstrated the utility of co-expression in understanding fungal pathogen biology [95,96]. However, these studies were performed in ascomycetes, which are closer to the model yeast *S. cerevisiae,* and thus there is much that can be inferred from orthology in these organisms. For the basidiomycete *C. neoformans,* there are more genes without clear orthologs and gene annotation information, making the need for computational predictions more urgent. Here, we leveraged publicly available RNAseq data from the *C. neoformans* research community to build a robust co-expression network for *C. neoformans.* We demonstrate that co-expression can predict gene function, both retrospectively as in the case of capsule and ergosterol biosynthesis, and prospectively, as in the case of DNA damage response proteins. Through co-expression, we were able to identify functions for 11 previously uncharacterized genes, including 6 that are specific to the Tremellales family. Overall, this demonstrates the utility of co-expression for predicting gene function in *C. neoformans*.

Co-expression network generation has been performed using multiple methods. We chose to use spearman rank correlation because it is relatively robust and interpretable, due its simplicity, and is among the top-performing co-expression methods in a recent benchmark [97]. We normalized expression scores based on Counts adjustment with TMM Factors (CTF) and Counts adjustment with Upper quartile Factors (CUF) normalizations. As an alternate to our approach, there is SNAIL, a method based on smooth quantile normalization aimed at reducing spurious associations [97]. Future refinements of the CryptoCEN network could use these alternate methods.

Importantly, CryptoCEN complements the current gene function annotation pipelines for *C. neoformans* by adding information not already captured from these databases. CryptoNet is based on a Bayesian integration of large-scale genomic and proteomic datasets, and this approach was effective at identifying genes involved in core virulence and drug resistance phenotypes [20]. Combining CryptoCEN with CryptoNET increased the retrospective predictive accuracy, and for a specific capsule network, we observed complementary connections.

A persistent limitation of CryptoCEN, however, is that without initial annotations, we cannot propagate information across the network. In *C. neoformans,* due to the high number of genes without annotation, there are many instances in which there is no signature or annotation in the entire network. This is especially the case for *Cryptococcus* or Basidiomycete-specific genes. For example, CNAG_00080 is a hypothetical protein that is *Cryptococcus-*specific; however, it is lowly expressed, and the top co-expression partner has a score of only 0.78. Only 5 of the top 50 co-expressed partners have any annotation at all, and the others are all hypothetical or unspecified product. As another example, CNAG_00465 is highly expressed basidiomycete-specific gene with many co-expressed partners; however, all the partners are only annotated as hypothetical proteins and there is no GO term enrichment. In these cases, the co-expression network has no information that can be used to predict gene function for these unknown proteins. In the future, functional genomic screens of mutant libraries will be critical for building the baseline information needed to predict gene function. For this reason, we focused on extending our information about known processes in *C. neoformans*; the DNA damage and response pathway provided a clear opportunity to identify novel proteins involved in a core biological process. Moreover, DNA damage and response processes in pathogens are thought to be potential targets of therapeutic potential [21]. Future work will include mechanistic studies to determine how these specific proteins contribute to EMS resistance or hypersensitivity.

## Supplemental Figures and Tables

**Supplemental Figure 1:**
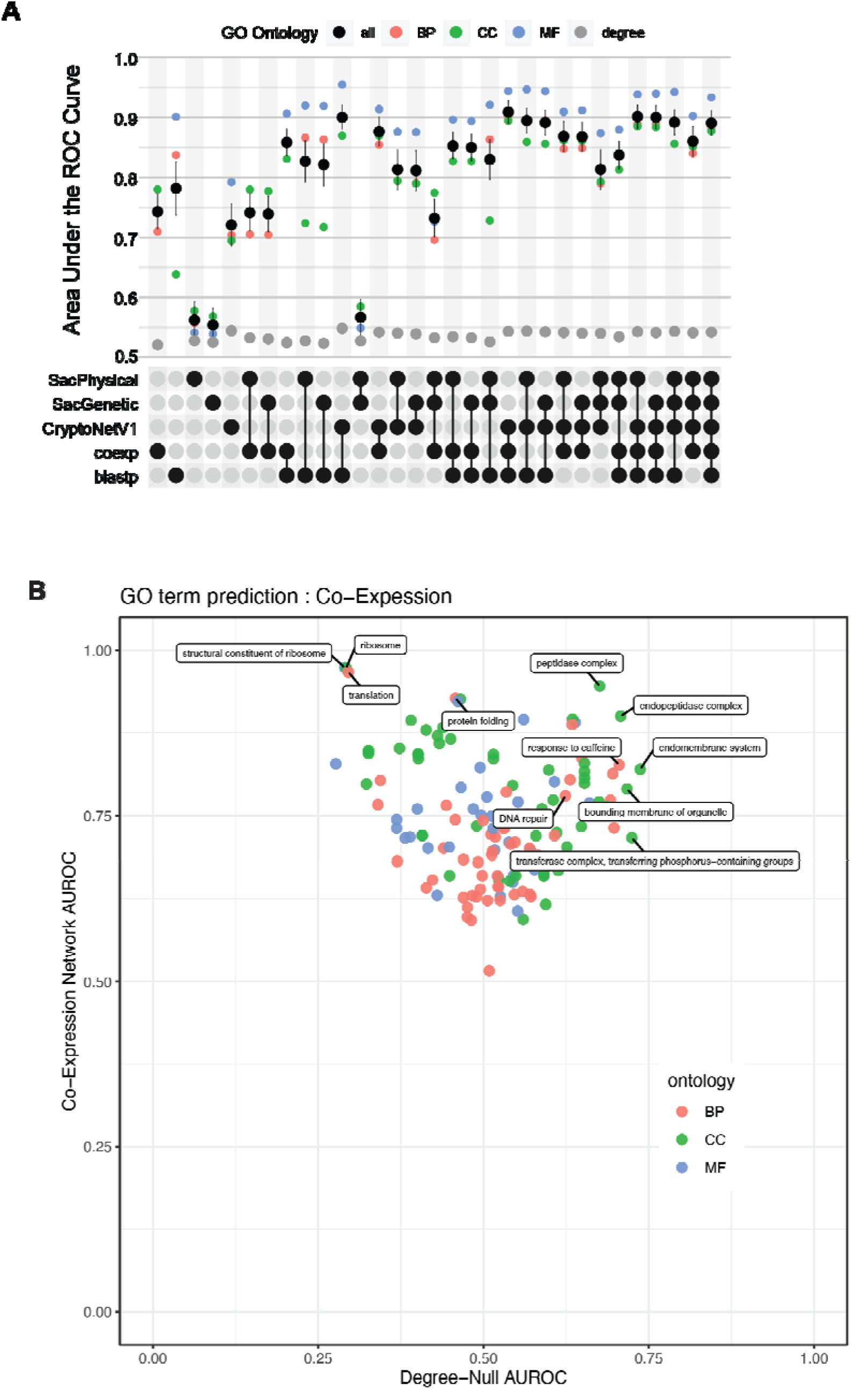
Retrospective predictive accuracy of CryptoCEN. A. UpSet plot of the retrospective prediction accuracy, as determined by the neighbor voting guilt-by-association (GBA) area under the ROC curve (AUROC). AUROCs values range between 0.5 for random predictor and 1 for a perfect predictor. As data sources are combined, the prediction accuracy increases. Each annotated GO term is colored by ontology biological process (BP), cellular component (CC), or molecular function (MF). **B.** Analysis of the centrality of specific GO terms to the co-expression network in *C. neoformans.* All the terms are above the diagonal, meaning that using the network structure increases the predictive accuracy above using each node in the network.

**Supplemental Figure 2:**
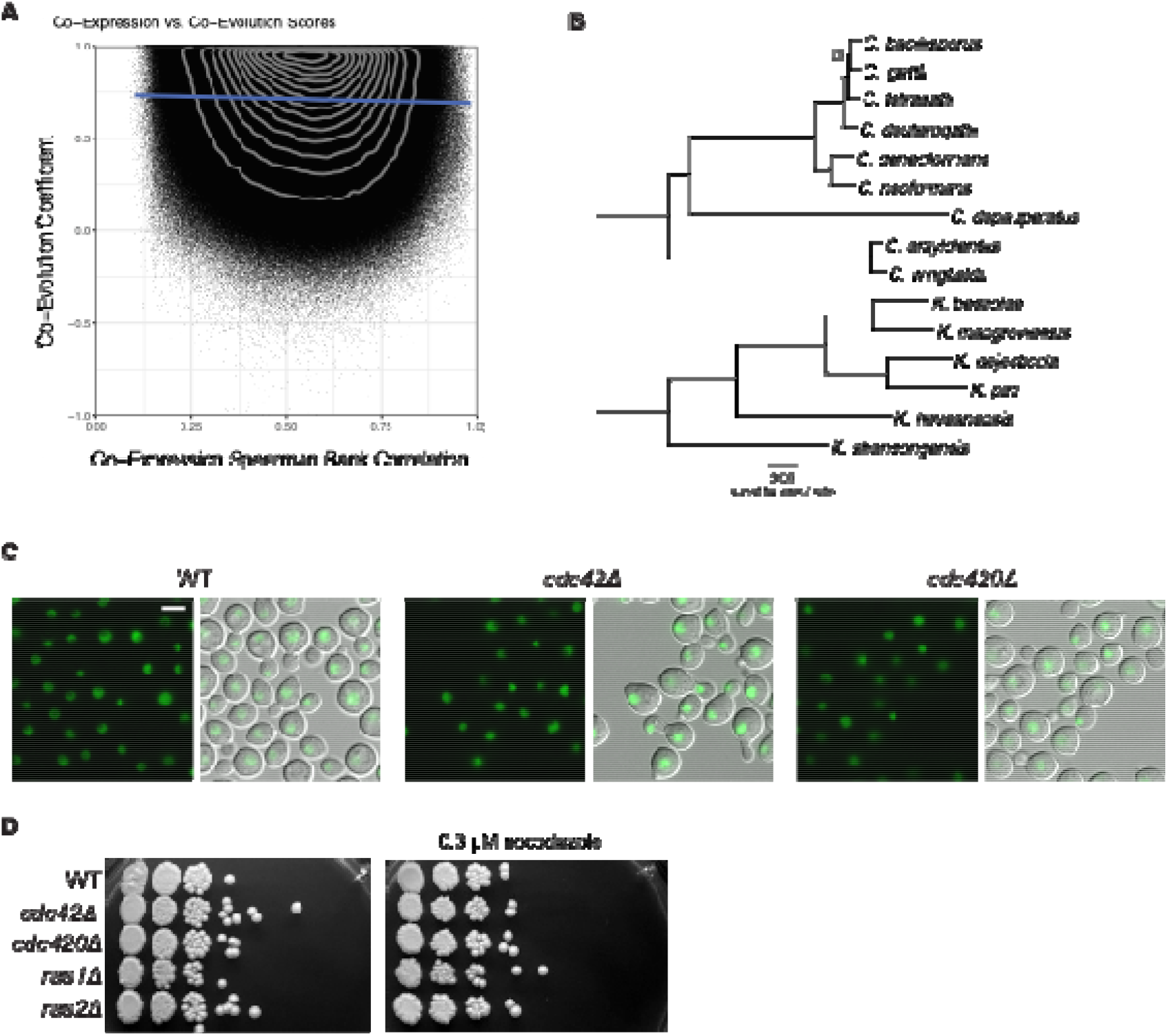
Evolutionary constraints on co-expression. **A)** Co-expression vs. co-evolution scores over the 5,264 overlapping gene sets does not show a positive correlation (correlation coefficient is −0.001). B) Phylogeny used for co-evolution. Tips indicate each species, scale bar indicates 0.05 substitutions/site. C) There is no difference in histone localization between the strains. Histones were marked by fusion of H4 with GFP. Cells were incubated in liquid YPD at 30 °C before imaging. Images taken at 40X magnification, scale = 5 microns. D) There is no difference in nocodazole sensitivity between the *cdc42*Δ or *cdc420*Δ mutant strains. The indicated strains were grown overnight at 30°C in liquid YPD medium and then serially diluted onto YPD or plates containing 0.3 µM nocodazole and incubated for 2 days before imaging.

**Supplemental Figure 3:**
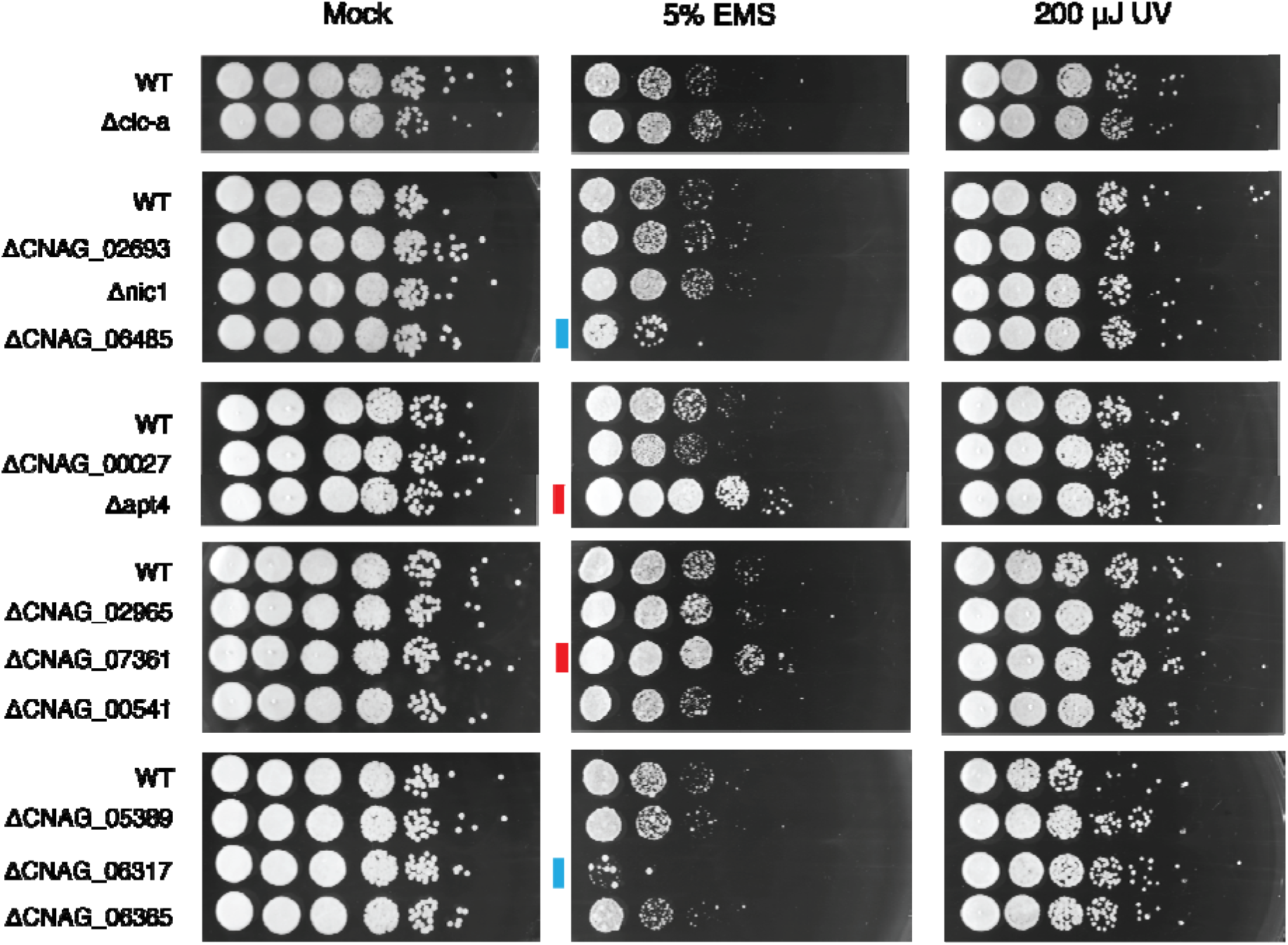
Matched controls do not show enrichment for DNA repair. A) Analysis of matched controls for phenotypes on DNA damaging agents. The indicated strains were grown overnight at 30°C in liquid YPD medium, and the 10-fold serially diluted cells were spotted onto YPD agar. For UV damage, the plates were immediately subjected to 200 µJ UV. For EMS, the cells were incubated in 5% EMS for 1 hr before serial dilution and plating. The plates were incubated at 30°C and imaged after 2 days.

**Supplemental Figure 4:**
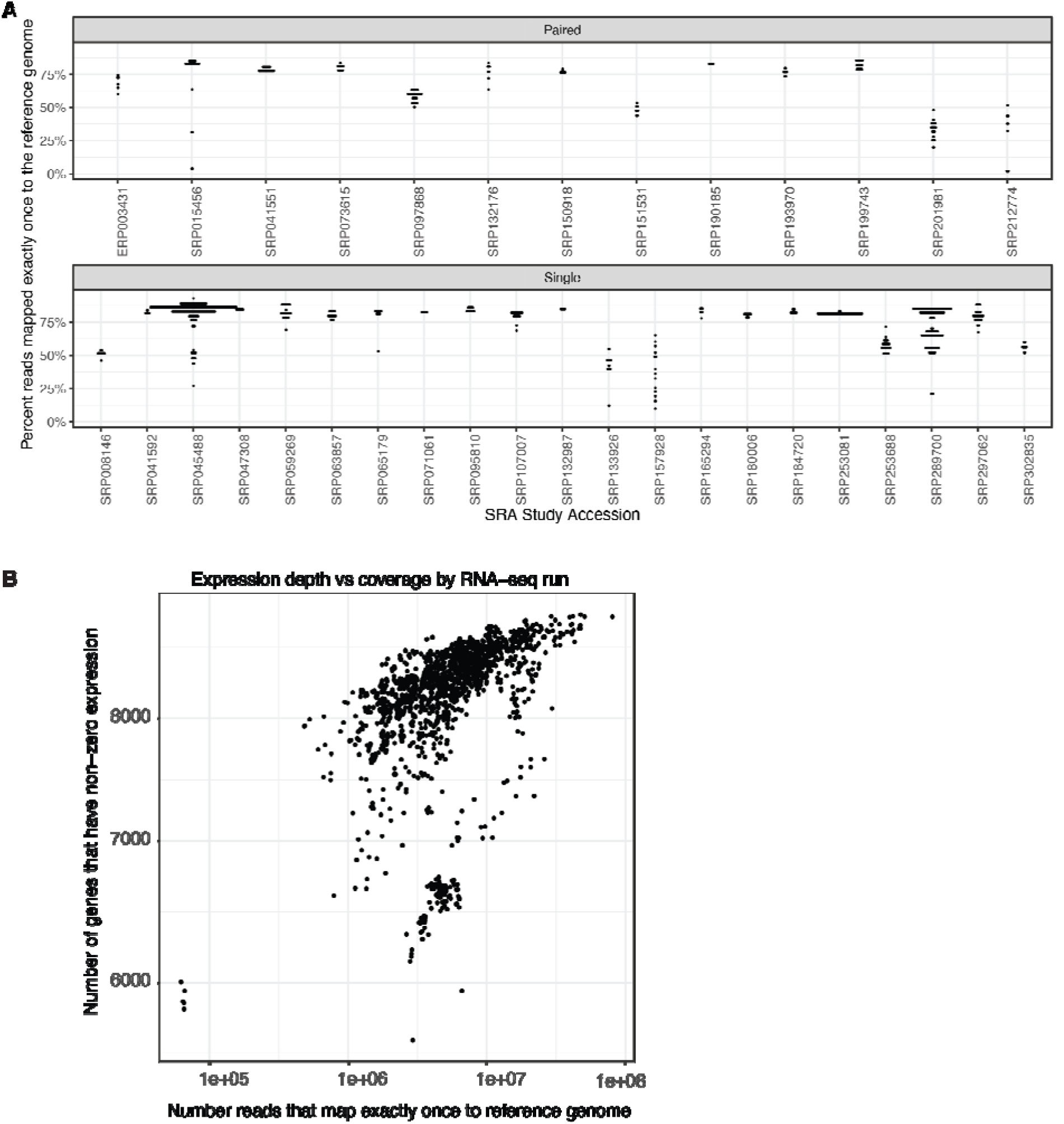
Mapping *C. neoformans* RNAseq runs. A. A total of 1,523 RNAseq runs from 34 identified studies are scatter-plotted as the number of genes with nonzero expression versus the fraction transcripts that map exactly once. B. The RNAseq runs that have nonzero expression for at least half of the genes which are used to construct the CryptoCEN network.

**Supplemental Figure 5:**
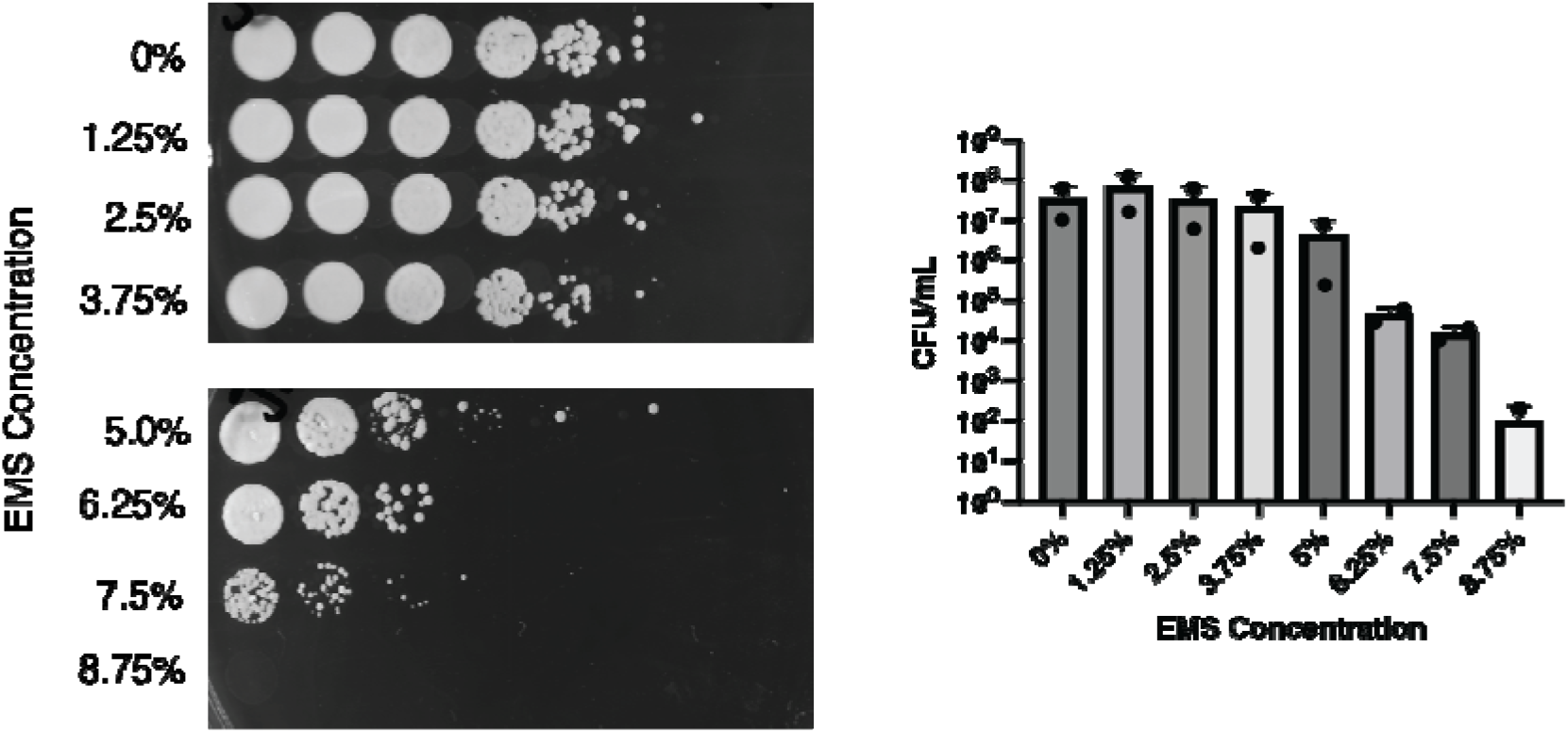
Establishing concentrations for EMS treatment. A) H99 wild type cells were incubated with the indicated concentrations of EMS for 1 hr before serial dilution and plating onto YPD. The plates were incubated at 30°C and imaged after 2 days. CFU/mL was calculated from the serial dilutions.

**Supplemental Table 1: Studies included in CryptoCEN**

Tab 1) RNAseq studies

Tab 2) RNASeq runs

**Supplemental Table 2: cluster analysis**

**Supplemental Table 3: Saccharomyces complexes**

Tab 1) summary

Tab 2) pairwise

**Supplemental Table 4: orthogroups**

**Supplemental Table 5: case studies**

Tab 1) capsule co-expression values

Tab 2) ergosterol co-expression values

**Supplemental Table 6: DNA damage**

Tab 1) co-expression

Tab 2) phenotype summary

Tab 3) reciprocal co-expression

**Supplemental Data 1: Full co-expression matrix**

**Supplemental Data 2: Full co-evolution matrix**

Supplemental data can be found at: DOI: 10.6084/m9.figshare.23979342

## Methods

### Collating *C. neoformans* RNAseq studies

Publicly available RNAseq studies were downloaded from SRA based on searches for *Cryptococcus neoformans* and filtered based on H99 or KN99 strains and mutant derivatives. For each study we collected the study accession, taxon, number of runs, study title and depositor. Using this information, where possible, we linked each study to a published journal article, and collected the experimental design and culture conditions for each strain.

### Mapping RNAseq reads

RNA expression was estimated by aligning reads to *Cryptococcus neoformans* H99 coding transcripts using release 49 from FungiDB downloaded on 8/05/2021 from https://fungidb.org/common/downloads/release-49/CneoformansH99/fasta/data/FungiDB-49_CneoformansH99_AnnotatedTranscripts.fasta. This release contains 9,189 ORFs defined by distinct unique cnag_id identifiers. Of these, 4 are labeled as pseudo genes and 838 are labeled as alternative splicing variants. Uncharacterized proteins were defined as non-pseudo gene, and the description does not contain “hypothetical protein” or “unspecified product”. SRA files were downloaded using prefetch and converted to FASTQ format using fastq-dump from the NCBI SRA-ToolKit package and then aligned using the RSEM package v1.2.31 [28] with bowtie2 [29] using the default settings. To assess sample quality, we measured the percent of genes with non-zero expression and number of reads that map uniquely to the C. *neoformans* genome. Of the 1,524 runs, the average percent genes with non-zero expression was 89%, with a minimum of 61%. The mean number of reads that mapped uniquely to the *C. neoformans* genome was 7.1M, and the minimum was 62k (SI Figure 4). For each study, the co-expression rank was estimated by the Spearman rank correlation coefficients of the FPKM values across all runs using in the study the EGAD R package (24) and then averaged across all studies to give the final network.

### Complementary networks

To build the sequence homology-based BlastP network, we used Protein-Protein BLAST 2.12.0+(BlastP) to compute sequence similarity between all C. *neoformans* open reading frames. We then ranked the bit-scores, and scores having identical values were given the average rank, then converted into a network using the build_weighted_network() function from the EGAD package.

Data for S. *cerevisiae* S288C was gathered from the Saccharomyces Genome Database (SGD) including the reference genome, all translated open reading frames, and annotations to the slim subset of the gene ontology using release 64-3-1 from 4/21/2021. S. *cerevisiae* genetic and physical interactor networks were built from data collected from BioGRID release 4.4.216, filtering for interactions with experimental system type of ‘genetic’ and ‘physical’ respectively, mapping to *C. neoformans* open reading frames by BlastP orthology, and building into networks using the build_binary_network() from EGAD package. The sparse binary networks were then extended defining edge weights as the inverse path length using the extend_network() function from the EGAD package.

We downloaded the CyproNetV1 from http://www.inetbio.org/cryptonet/ across all 14 data type specific networks, which contains 156,506 edges across 5,649 genes.

### Embedding of the Co-Exp network

Using the R monocle3 package, we pre-processed the gene by study co-expression matrix by PCA and then used UMAP to non-linearly reduce to 2 dimensions with parameters a = 30, b = 0.8, and default parameters otherwise. To cluster, we used Louvain clustering with parameters k=30, num_iter = 15, resolution = 0.001. We then plotted the embedding coloring by cluster using ggplot2. To assign functional annotations to each cluster we used GO term enrichment through FungiDB, focusing on the biological process information.

### Retrospective function AUROC calculations

13,920 Functional annotations for C. *neoformans* were downloaded from FungiDB release 49 and mapped to GO ontology terms using the GO.db R package gathered on 8/19/2021. Annotations with the NOT qualifier were excluded, and the remaining annotations were propagated along ‘isa’ and ‘part of’ relationships in the GO ontology, yielding 23,863 annotations. Then, to facilitate guilt-by-association gene function prediction, terms with more than 1000 or less than 20 annotations were excluded, yielding 14,215 annotations across 3,421 open reading frames for 145 terms with an average of 4.6 annotations per open reading frame and 98 annotations per go_id.

### Co-expression of *S. cerevisiae* Complexes analysis

616 *S. cerevisiae* complexes were downloaded from EBI Complex Portal (v2021-10-13)[98]. Members with identifiers that began with ‘URS-’ (RNA from rnacentral.org id), ‘EBI-‘ (mRNA), ‘CHEBI-‘ (small molecule), ‘CPX-‘ (other complex) or ‘P12294’ (mitochondrially encoded), yielding 1,948 proteins from S. *cerevisiae* strain S288C (vR64-2-1) that were mapped to genes in the Saccharomyces Genome Database[99] using UniProtKB[100]. Of these genes, 1,093 mapped 1-to-1 to *C. neoformans* genes via uniport accession to gene identifier and OrthoMCL (v6.7)[101] based orthology with an additional 211 genes via BlastP based orthology, yielding 1,304 *C. neoformans* genes predicted to participate in 558 complexes (**Table S3**). Among the *C. neoformans* orthologs in these complexes, there are 13,950 distinct co-complex associations. 51% C. *neoformans* co-complex associations are within 17 complexes with “ribosomal” or “ribonucleoprotein” in their name.

### Orthology with S. *cerevisiae*

Pairwise sequence-based orthology using BlastP was defined by reciprocal best hits and having E-values less than 1e-5 in both directions yielding 2,248 associations. <u>Ortholog based on</u> <u>sequence-based clustering using OrthoMCL release 6.7 was defined by orthogroups containing</u> <u>exactly one member from C. *neoformans* and one from S. *cerevisiae*, yielding 2,274</u> <u>associations.</u>

### Co-evolution coefficient determination

To identify gene pairs with signatures of co-evolution, which refers to the covariation of the relative evolutionary rates in two genes across speciation events [102], we used the Covarying Evolutionary Rates (CovER) function in PhyKIT, v1.11.10 [103]. This analysis requires phylogenetic trees of single-copy orthologs from a panel of species. To generate this data, 15 genomes from *Cryptococcus* and the sister genus *Kwoniella* were obtained from NCBI, which spans all publicly available annotated genomes at the time of downloading (07/2022). Orthology was inferred using OrthoFinder, v2.3.8 [104], an algorithm that conducts graph-based clustering of global sequence similarity values calculated using DIAMOND, v2.0.13.151 [105]. Orthology was inferred among protein sequences using an inflation value of 1.8 resulting in 3,828 single-copy orthologs.

To obtain additional groups of orthologous genes for coevolutionary coefficient determination, the OrthoSNAP pipeline, which identifies single-copy orthologs nested within larger gene families [106], was used. To do so, phylogenetic trees were inferred from protein sequences of multi-copy orthogroups with at least eight taxa using IQ-TREE, v2.0.6 [107], with 1,000 ultrafast bootstrap approximations [108]; multi-copy orthogroups were first aligned using MAFFT, v7.402 [109], with the auto parameter, and trimmed using ClipKIT, v1.3.0 [110], with default parameters. The resulting phylogenies were used as input into OrthoSNAP, v1.0.0, which resulted in 1,630 additional single-copy orthologs (5,458 total single-copy orthologs).

The resulting single-copy orthologs were concatenated into a supermatrix using the “create_concat” function in PhyKIT, v1.11.10 [103] and used for species tree estimation using IQ-TREE 2 (best fitting substitution model: JTT+F+I+G4). For each single-copy ortholog, branch lengths were inferred along the species tree using IQ-TREE 2. For every pairwise combination, the coevolutionary coefficient was calculated using the “cover” function in PhyKIT, v1.11.10 [103]. In brief, PhyKIT identifies pairs of coevolving genes by first accounting for confounding variables like time since divergence and mutation rate by correcting single-copy ortholog branch lengths with the corresponding branch length in the species tree; the resulting values are Z-transformed and used for Pearson correlation analysis, representing a quantitative measure of gene-gene coevolution.

### Paralog identification

Orthology information from the previous section was used to determine pairs of paralogs. 6,641 ortholog groups were mapped over 6,975 C. *neoformans* genes. Among these groups, we identified 1,056 paralog associations across 550 genes.

### Strain construction

The following *C. neoformans* strains were generated by mating crosses which consisted of co-culturing strains of opposite mating type on MS medium for 7-days [111]. Recombinant progeny were isolated from the mating mixture by random spore dissection and analyzed for genotype, phenotype, and fluorescence. A mating cross between MATa *cdc42*Δ*::nat* (ERB010) and a MATα wild type strain expressing GFP-tagged alpha tubulin (*TUB1a*; CBN242) generated *cdc42*Δ*::nat* + *GFP-TUB1a* (CBN587). Similarly, a mating cross between MATa *cdc420*Δ*::neo* (ERB007) and CBN242 generated *cdc420*Δ*::neo* + *GFP-TUB1a* (CBN589). Mating crosses of strains ERB010 and ERB007 with a MATα wild type strain expressing a GFP-tagged Histone H4 (CNV108) generated *cdc42*Δ*::nat* + *GFP-H4* (CBN594) and *cdc420*Δ*::neo* + *GFP-H4* (CBN593).

### Nocodazole sensitivity assay

To test sensitivity, each strain was 2-fold serially diluted at 5 µL spots of each dilution were plated onto YPD with or without the indicated concentration of nocodazole. Plates were incubated for 3 days before imaging.

<u>Histone / tubulin localization assay:</u> The following strains were incubated to mid-logarithmic phase with shaking (200 rpm) in YPD medium at either 30°C or 37°C, with or without the addition of 0.125 μM nocodazole: strains CNV108 (WT + Gfp-Histone H4), CBN242 (WT + Gfp-Tub1), CBN593(*cdc420*Δ + Gfp-Histone H4), and CBN589 (*cdc420*Δ + Gfp-Tub1). Nuclear size and division (noted by Gfp-H4) and tubulin filaments (Gfp-Tub1) were assessed using a Zeiss Axio Imager A1 microscope equipped with an Axio-Cam MRmdigital camera. Cells were imaged by DIC and with eGFP filter. Identical exposure times were used to image all cells. Fiji software [112] was used to process images.

### DNA Damage Response Assay

EMS and UV mutagenesis assays were based on the protocol described by Winston [113]. Overnight cultures of *C. neoformans* were normalized to an OD_600_ of 1. Cultures were then centrifuged in 1 mL aliquots and washed and resuspended in 0.1 M sodium phosphate buffer (pH 7.4) to remove excess YPD. For EMS mutagenesis, cells were then transferred to 15 mL conical tubes where either 5% ethyl methanesulfonate or sodium phosphate buffer was added. Dose of EMS was chosen after dose-response assays **(Figure S6).** Final volume in each tube was adjusted to 2 mL and cells were incubated at 30°C for 1 hour while shaking. After incubation, all samples were pelleted and washed with a 5% sodium thiosulfate solution to inactivate the EMS. Following this wash, cells pellets were resuspended in 1 mL of sterile water and diluted 10-fold. 5 µL of each dilution were spotted onto YPD plates and incubated for 48 hours before imaging. For UV mutagenesis, mock-treated samples were spotted onto YPD, but exposed to 200 µJ of UV radiation using a UV-Stratalinker before incubation at 30°C for 2 days before imaging on a BioRad gel dock.

## Acknowledgements

We thank Vikas Yadav for thoughtful comments on kinetochores.

## Abbreviations

RNAseq: RNA sequencing
GO: gene ontology
BP: biological process
CC: cellular component
MF: molecular function
SRA: sequence read archives
FPKM: fragments per kilobase of transcript per million mapped reads
GBA: guilt-by-association
AUROC: area under the receiver operating characteristic curve
ATRIP: ATR-interacting protein
GEF: guanidine exchange factor
LOESS: locally estimated scatterplot scattering
EMS: ethyl methanesulfonate

