## Supplementary figures and images for "CryptoCEN: A Co-Expression Network for *Cryptococcus neoformans* reveals novel proteins involved in DNA damage repair"

### Supplemental Figure 1

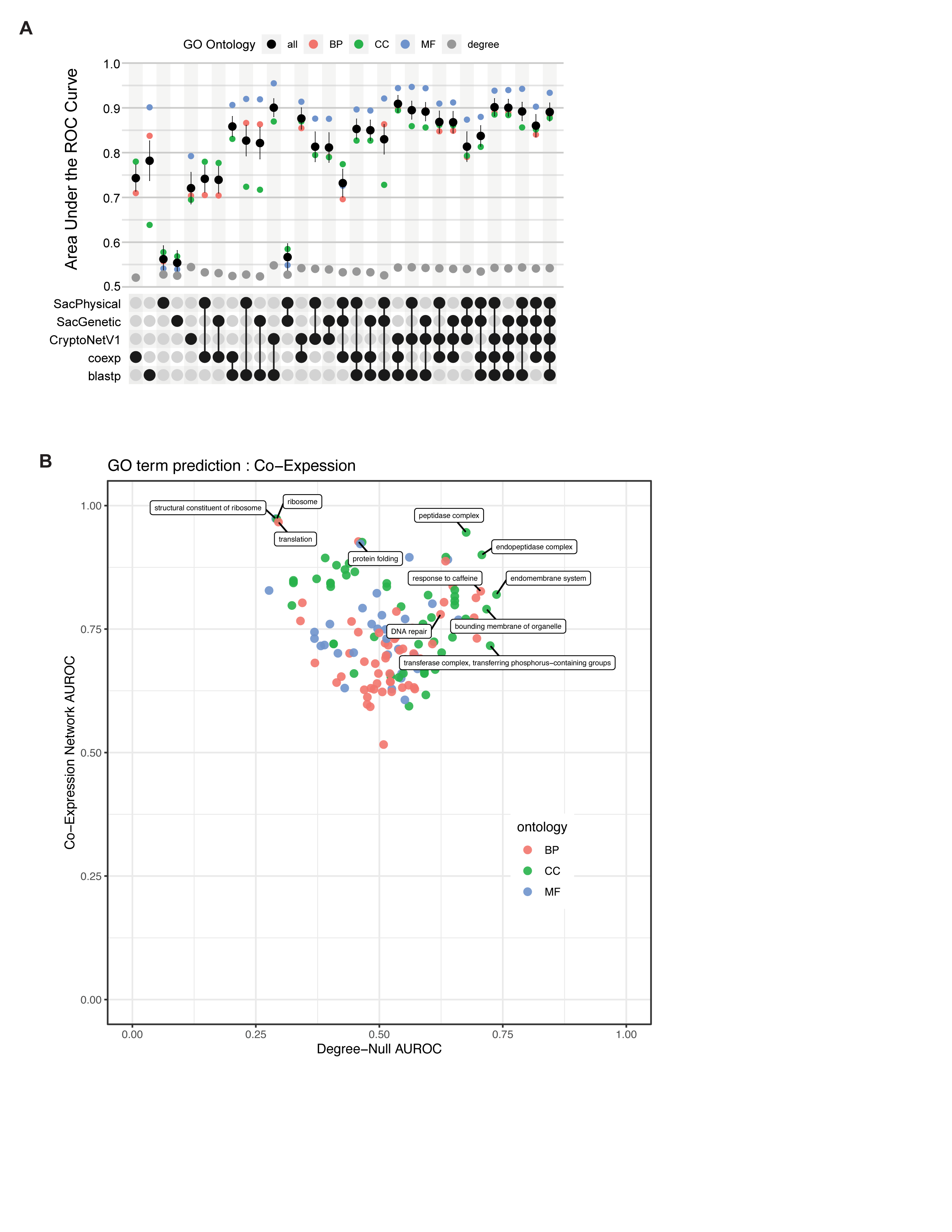

### Supplemental Figure 2

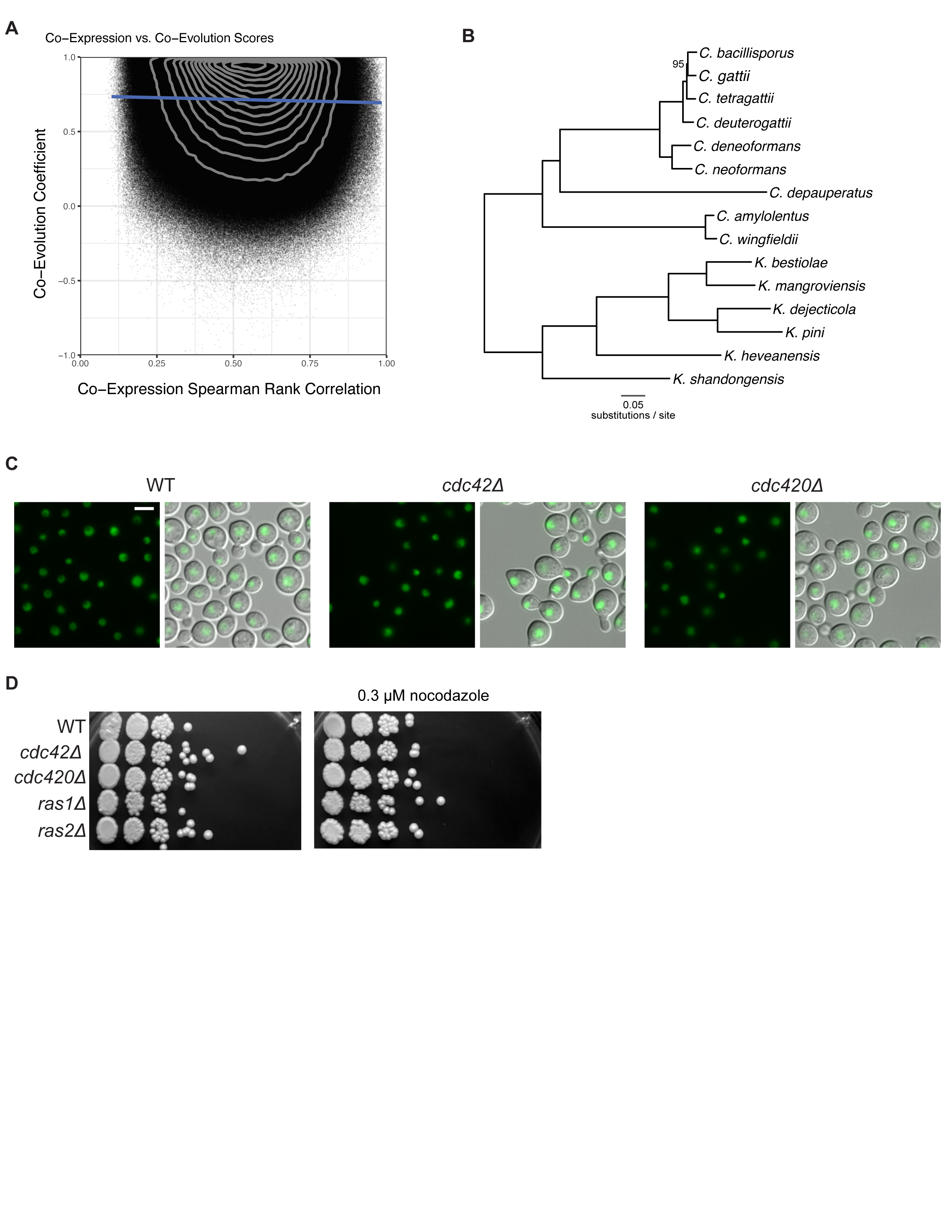

### Supplemental Figure 3

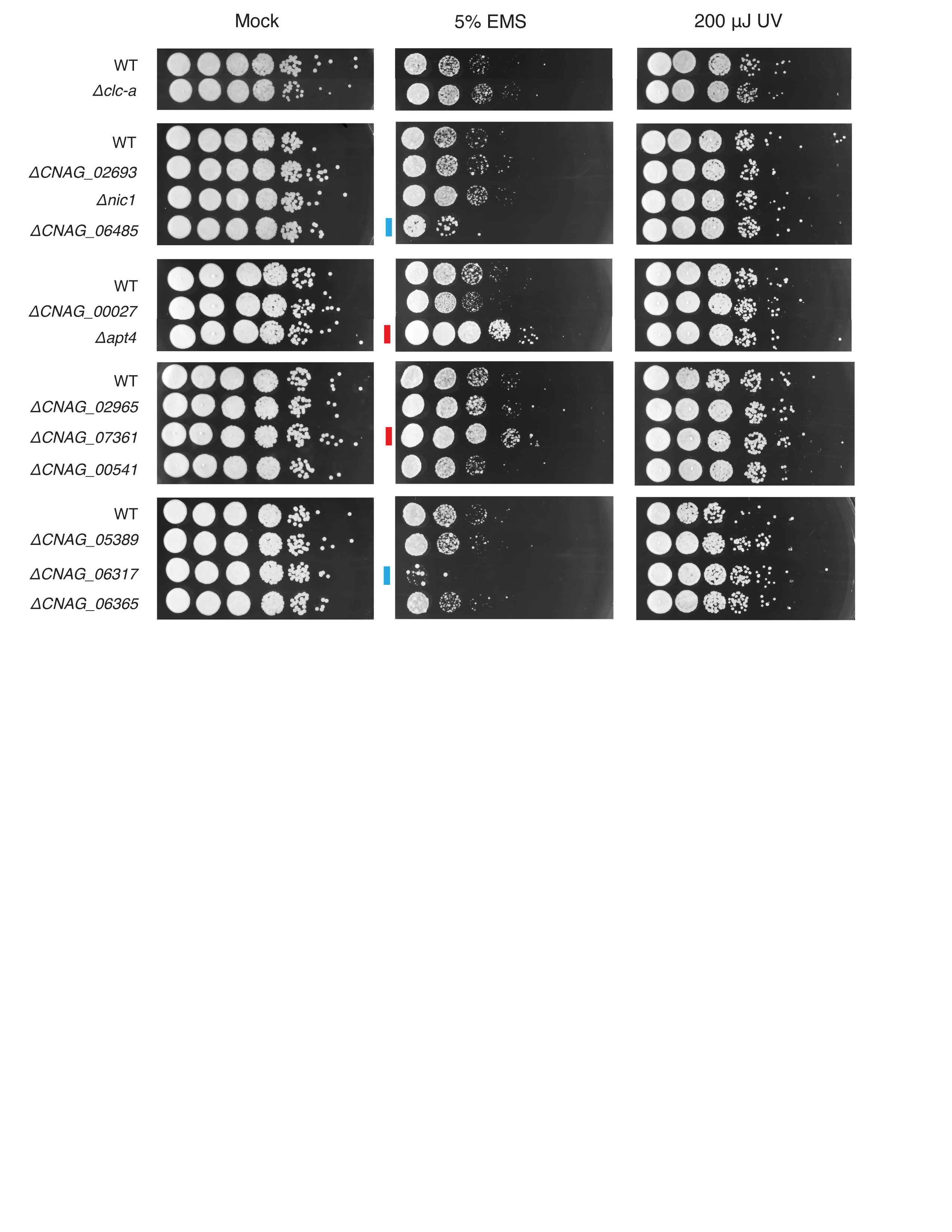

### Supplemental Figure 4

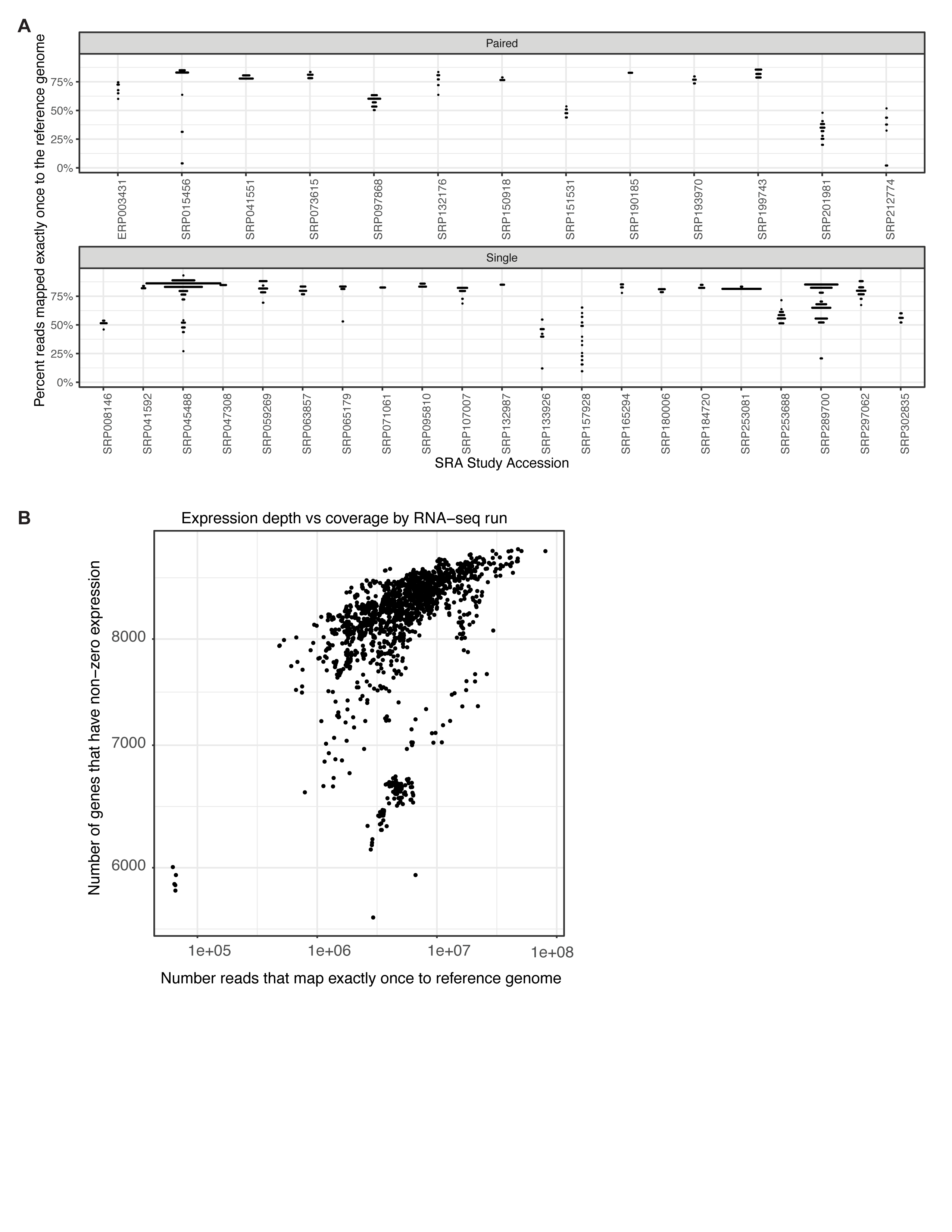

### Supplemental Figure 5

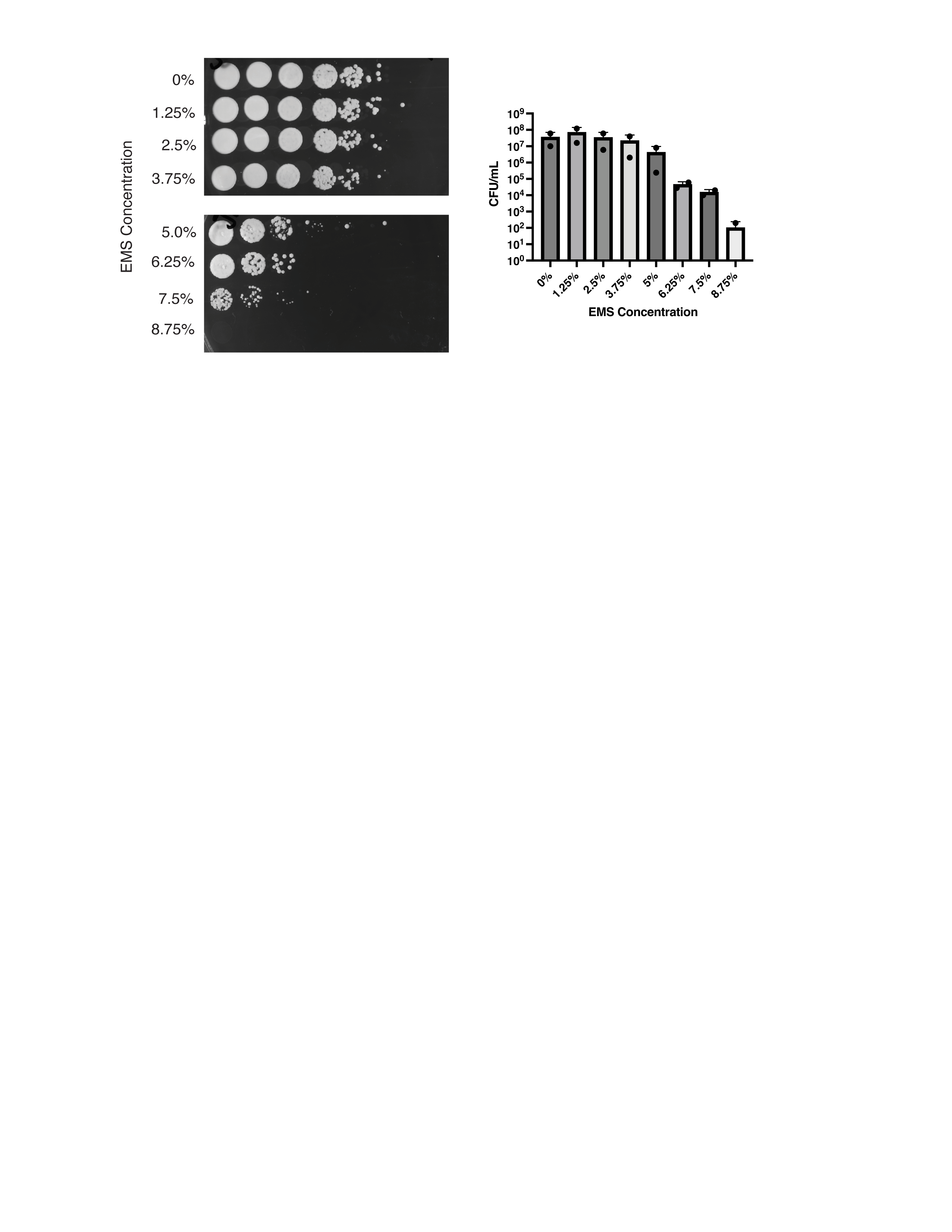
